# Mapping and Harmonization of CVX vaccine terms to the Vaccine Ontology

**DOI:** 10.1101/2025.07.15.664501

**Authors:** Yuanyi Pan, Warren Manuel, Rashmie Abeysinghe, Jie Zheng, Alexander Davydov, Qi Yang, Asiyah Yu Lin, Licong Cui, Yongqun Oliver He

## Abstract

**Background:** With many vaccines developed and used, it is critical to standardize vaccine information. The OHDSI OMOP Common Data Model (CDM), widely used to support EHR data integration and analysis, leverages CVX, RxNorm, and RxNorm Extension codes to standardize vaccine-related records. However, these terminologies lack robust semantic relations, making the vaccine classification ineffective in OMOP CDM. To address this issue, our OHDSI Vaccine Vocabulary Working Group proposes to use the Vaccine Ontology (VO) to map these standards and build up its own semantic relations. As a first study of the work, we performed the mapping and alignment of the Vaccine Administered (CVX) codes with the VO using a combination of semi-automatic and manual mapping methods.

**Results:** A total of 273 CVX terms were first collected and classified. A high-level VO design pattern and an exact one-to-one mapping strategy were developed to guide the CVX-to-VO term mapping. To facilitate the manual mapping and harmonization process, we also developed and evaluated three semi-automated mapping approaches utilizing lexical and semantic information of vaccine concepts to map CVX to VO. These approaches suggested candidate VO mappings for CVX terms and also indicated CVX terms that were unmappable to VO and required new term additions to VO. The application of the best approach to the 2022-10-05 release of VO achieved an accuracy of 85.55% for its suggestions. The suggestions made by the semi-automated approaches were taken into account to further enhance the mappings, which led to our eventual mapping of all CVX terms to the latest version of VO. We innovatively proposed the inclusion of the ‘passive vaccine’ branch in VO, which includes 24 immunoglobulins and antitoxins from CVX as passive vaccines. A specific CVX-VO OWL file was developed and added to the VO GitHub. Use case queries were developed to demonstrate its support for computer-assisted queries of vaccine groups based on CVX-VO hierarchies.

**Conclusion:** All CVX terms were mapped to the VO using our combined semi-automatic and manual mapping methods. The mapped results enhanced semantic vaccine classification, providing a basis for further OMOP vaccine classification and EHR data analysis.

## 1. Introduction

Vaccines have played an important role in fighting against infectious diseases such as COVID-19. In recent decades, a large amount of vaccine-related electronic health records (EHR) have been produced. The Observational Medical Outcomes Partnership Common Data Model (OMOP CDM) is a widely used open community data standard model that has significantly promoted the standardization of the structure and content of observational data, including vaccine data, and to enable efficient data analyses^1^. A core innovative part of the Observational Health Data Sciences and Informatics (OHDSI) OMOP system is its reuse of many standard vocabularies, including terminologies and ontologies to standardize all kinds of EHR data, leading to the ability of integrating the large and often heterogeneous EHR data from different data resources.

The current OMOP approach of handling vaccine vocabularies has its advantages but also has drawbacks. Currently, OMOP CDM and its associated vocabularies (e.g., CDC Vaccine Administered (CVX), RxNorm, and SNOMED-CT) include a variety of vaccine-related terms. However, these vaccine vocabularies have different coverages and use different design patterns and representation styles^1^. As a result, the vaccine terms in these vocabularies could not be easily mapped and integrated. Previous studies have further systematically analyzed the OMOP vaccine vocabularies and found a number of mapping issues such as erroneous mapping and missing mapping, which may lead to discrepancy of source data integration^2,3^.

To address the issues with the OMOP vaccine vocabulary, our current OHDSI vaccine vocabulary working group has proposed to use the Vaccine Ontology (VO) as a way to map, align, and build strong semantic relations among vaccine terms. The VO^4^ is a community-based biomedical ontology in the vaccine domain that provides a systematic ontological representation of all vaccines at different stages (including the licensed stage, clinical trials, and research stage), the vaccine components (e.g., vaccine antigen, vaccine adjuvants), vaccine responses, and their internal relations. As a reference ontology in the domain of vaccinology, the VO has been accepted as a member of the Open Biological and Biomedical Ontology (OBO) Foundry ontology library^5^. The OHDSI vaccine working group believes that the VO has the ability to address the issues of CVX and RxNORM including their lack of semantic hierarchical structures and needed relations (e.g., vaccine types and targeted diseases and pathogens), and the working group has aimed to explore the possibility and feasibility of adding VO as a vocabulary to address these issues. VO has included over 4,000 vaccines but it does not include more of the CVX and RxNorm terms. Using the VO would support interoperability, ensure consistent communication across different healthcare systems, and facilitate effective ontology-based data analysis.

Our current focus is to perform a comprehensive mapping between CVX and VO. The CVX is a product of the National Center of Immunization and Respiratory Diseases (NCIRD) in the Centers for Disease Control and Prevention (CDC)^6^. As part of the Health Level Seven International (HL7), CVX has been widely used for EHR data integration and analysis. For example, the Vaccine Adverse Event Analysis System (VAERS) uses CVX to classify the vaccines^7^. CVX has also been incorporated into the OMOP vocabulary system^8^. However, although CVX is relatively small compared to RxNorm, the relations among CVX terms are still very complex and how they are associated with vaccine terms in other vocabularies such as RxNorm is unclear. Therefore, a more systematic semantic modeling and mapping is desired.

The mapping of external vocabulary terms to VO can be time-consuming and labor-intensive if it is performed entirely manually. The experts must comprehensively go through both VO and the external vocabulary to identify matches to ensure accuracy and consistency. Automated or semi-automated approaches can support and complement the manual process by increasing efficiency and reducing the overall workload^9^. In this paper, we explore three semi-automated mapping approaches that suggest a number of candidate VO mappings for an external vocabulary term. If the VO does not contain an appropriate term to match an external vocabulary term, then an appropriate method is required to determine that this is an unmappable situation requiring the addition of a new concept to VO. The curators review the suggestions made by the method and make changes to VO where appropriate, thereby significantly reducing their manual effort.

This manuscript reports complementary efforts to achieve the common theme of CVX-to-VO mapping and analysis. Our complementary efforts include the high level design for the CVX-to-VO mapping, semi-automatic mapping that combines automatic mapping algorithm development and manual evaluation, and demonstrations of how such mapping can be helpful.

## 2. Methods

### 2.1. Collection and classification of CVX terms

The CVX vaccine terms were extracted from the CDC website^6^, May 22, 2024. Other publicly available references were also systematically surveyed^10^. Since existing CVX vaccine terms are usually provided as a single list, we classified them primarily based on their target disease or its causing pathogen groups.

### 2.2. Initial stage of CVX-to-VO mapping and VO updating

A combination of top-down and bottom-up methods was used. Specifically, our top-down strategy was to evaluate the existing high-level VO hierarchical design and then propose an enhancement of the VO high-level design in order to eventually incorporate all the CVO terms to VO. For example, the original VO did not have a solid strategy for incorporating many immunoglobulin or antibodies against different diseases, then we updated our VO to include a ‘passive vaccine’ branch so that we were able to include all the disease-specific immunoglobulin terms in CVX to the new branch. To support CVX-specific annotations in VO, three specific annotation properties, including ‘*CVX code*’, ‘*CVX full name*’, and ‘*CVX short description*’ were added to VO to represent the corresponding CVX information.

A specific mapping directionality was also designed. Three options were identified: a hierarchy (the mapped VO term is always the same or more general to the CVX term), polyhierarchy (one-to-many term mapping allowed), or one-to-one exact mapping. Specifically, our manual mapping targets for one-to-one exact mapping. It means that for any single CVX term, we will have one and only one VO term that maps to exactly the CVX term. Meanwhile, we might add new higher level terms in order to classify or group a set of CVX terms with the defined feature such as the function against influenza viral infection. Our mapping does not allow downhill mapping. For example, we do not use a specific vaccine with brand name such as ‘*M-M-R II’* (VO_0000069) to map to a CVX term such as ‘*measles, mumps and rubella virus vaccine*’ (CVX code: 03). In this case, we use the VO term ‘*Measles-Mumps-Rubella vaccine*’ (VO_0000731) to map the CVX term, which is the exact match and one-to-one mapping. When an exact one-to-one mapping does not exist in VO, a broader mapping that maps the term to its immediate parent term by an automatic mapping algorithm is considered accurate. Such a broader mapping would help us to find the immediate parent term in VO and support later addition of a new term to VO.

For the bottom-up strategy, we started to select a small set of CVX terms for manual VO mapping. The VO source was obtained from VO GitHub^11^. Ontobee^12^ was used to query vaccine terms from the VO and related ontologies. A manual evaluation was performed for VO-CVX term mapping. For those CVX terms with no corresponding VO mapping, we generated new VO terms based on the standard VO design pattern^13^. The Protege OWL editor^14^ was used for manual editing, and the Ontorat tool^15^ was used for automatic addition of VO terms based on pre-defined ontology design patterns. To support the mapping process, an Excel file (freely accessible in the VO GitHub as detailed later in the manuscript) was developed to list individual CVX terms and associated information. The small set of manually annotated CVX terms were later provided to the automatic mapping group for their training and testing as detailed below.

### 2.3. Semi-automated approaches for CVX-VO mapping

There were two purposes for the development of the semi-automated approach: (1) automatically identifying candidate VO mappings for a given CVX term; and (2) automatically identifying candidate CVX concepts that do not have a mapped VO term, hence needing a new VO concept. The findings of the approaches were handed over to the curators for review and potential adoption to VO.

To begin this process, we first extracted vaccine terms from both CVX and VO. Of the 273 CVX terms present, 10 terms related to tuberculin skin tests and other miscellaneous non-vaccination records were not considered for mapping. In the same manner, to ensure that only relevant concepts would be present in the candidate pool within VO, only concepts under the sub-hierarchy “*vaccine*” (VO_0000001) were selected to be considered. In order to identify the candidate VO concepts for each CVX term, three similarity-based approaches were investigated: (1) Embedding-based similarity approach; (2) World-level similarity-based refinement; and (3) Large Language Model-based refinement.

If a method does not generate at least one candidate VO concept for a given CVX term, this particular CVX term is considered to be unmappable to VO. In such situations, we recommend adding a new concept to VO to encompass the information represented by the CVX term.

#### 2.3.1. Pre-processing

Prior to similarity calculation across all the methods, lexical information of terms was normalized via lowercase and ASCII conversion, and expansion of common vaccine-related abbreviations and trade names sourced from the CDC^16,17^. Further, common stopwords in the English language, punctuations, and the term ‘vaccine’ were removed from these terms. For VO, lexical information included the term labels as well as all synonyms available.

#### 2.3.2. Embedding-based similarity approach

This approach leveraged sentence encoders pre-trained on large volumes of biomedical data to obtain embeddings for concepts that are representative of their semantic meanings. For this purpose, MiniLM-L6-v2-pubmed-full^18^ a sentence encoder trained on PubMed documents was utilized. The higher-dimensional vector embeddings for each VO concept were then compared with each CVX term embeddings to identify the most similar terms. The similarities of these 384-dimensional embeddings were calculated using cosine similarity. To ensure that all lexical information within a VO concept was utilized, each synonym was considered individually and the best similarity score was used to represent the similarity between the CVX and VO concept. For example, when calculating the similarity score between concepts **CVX_130** ‘*Diphtheria, tetanus toxoids and acellular pertussis vaccine, and poliovirus vaccine, inactivated’* and **VO_0000742 ‘***Diphtheria-Tetanus-Pertussis-Poliovirus vaccine*’, both its label and two synonyms (1: *DTaP/IPV vaccine* and 2: *diphtheria and tetanus toxoids and acellular pertussis adsorbed and inactivated poliovirus vaccine*) were considered. Hence, for this concept pair, three similarity scores were calculated (0.8759, 0.5690, 0.9518, respectively), and the highest score of 0.9518 was retained as the similarity score between the two concepts. For a given CVX term, all the VO terms with a similarity score greater than 0.8 were considered as mapping candidates. This threshold was set based on experimentation.

#### 2.3.3. Word-level similarity-based refinement

In this approach, for each CVX term we first obtained the top 20 candidate VO matches by the embedding approach. These were further refined based on their word-level similarity to identify up to top-10 matches. The lexical information of these top-20 matches was tokenized prior to computing the word-level similarity.

For example, the concept **VO_0000743**: Diphtheria-Tetanus-Pertussis-Poliovirus-Hepatitis B vaccine and synonym DTaP/IPV/HepB vaccine after the preprocessing steps would yield the sets of tokens {*‘b’, ‘diphtheria’, ‘tetanus’, ‘pertussis’, ‘poliovirus’, ‘hepatitis’*} and {*‘b’*, *‘diphtheria’, ‘tetanus’, ‘acellular’, ‘pertussis’, ‘inactivated’, ‘poliovirus’, ‘hepatitis’*}. Note that the synonym usage and abbreviation expansion yielded the addition of the terms ‘acellular’ and ‘inactivated’. Therein, the lexical similarity of the 20 candidates obtained from the embedding method was computed via the Jaccard similarity^19^, where the coefficient is calculated as the intersection between tokens of the two concepts divided by the union of the two. For example, to calculate the Jaccard similarity between ***CVX_54***: ‘*adenovirus vaccine, type 4, live, oral*’ and ***VO_0004872***: *‘Adenovirus Type 4 Vaccine Live Oral Product‘;* the concept labels after preprocessing would yield the tokens; ***CVX_54:*** {*4, type, oral, adenovirus, live*}, and ***VO_0004872***: {*product, 4, type, oral, adenovirus, live*}. Where the intersection of tokens for the two concepts being {*4, type, oral, adenovirus, live*} (n=5) and the union being {*product, 4, type, oral, adenovirus, live*} (n=6). Hence, the Jaccard similarity between the two concepts would be ⅚ = 0.833.

#### 2.3.4. Large Language Model-based refinement

Similar to the word-level similarity-based refinement approach, we first obtained the top-20 candidate VO matches by the embedding method. These were further refined by leveraging the Large Language Model GPT-4o^20^. For this step, a prompt was engineered to identify up to 10 candidates out of the 20 that are either synonymous or more general. The developed prompt is shown in **Figure 1**. The prompt contains an introduction to the task, including the inputs and outputs, an example, and the CVX term and the candidate vaccine ontology term list (under “Begin with the following input” section).

**Figure 1.**
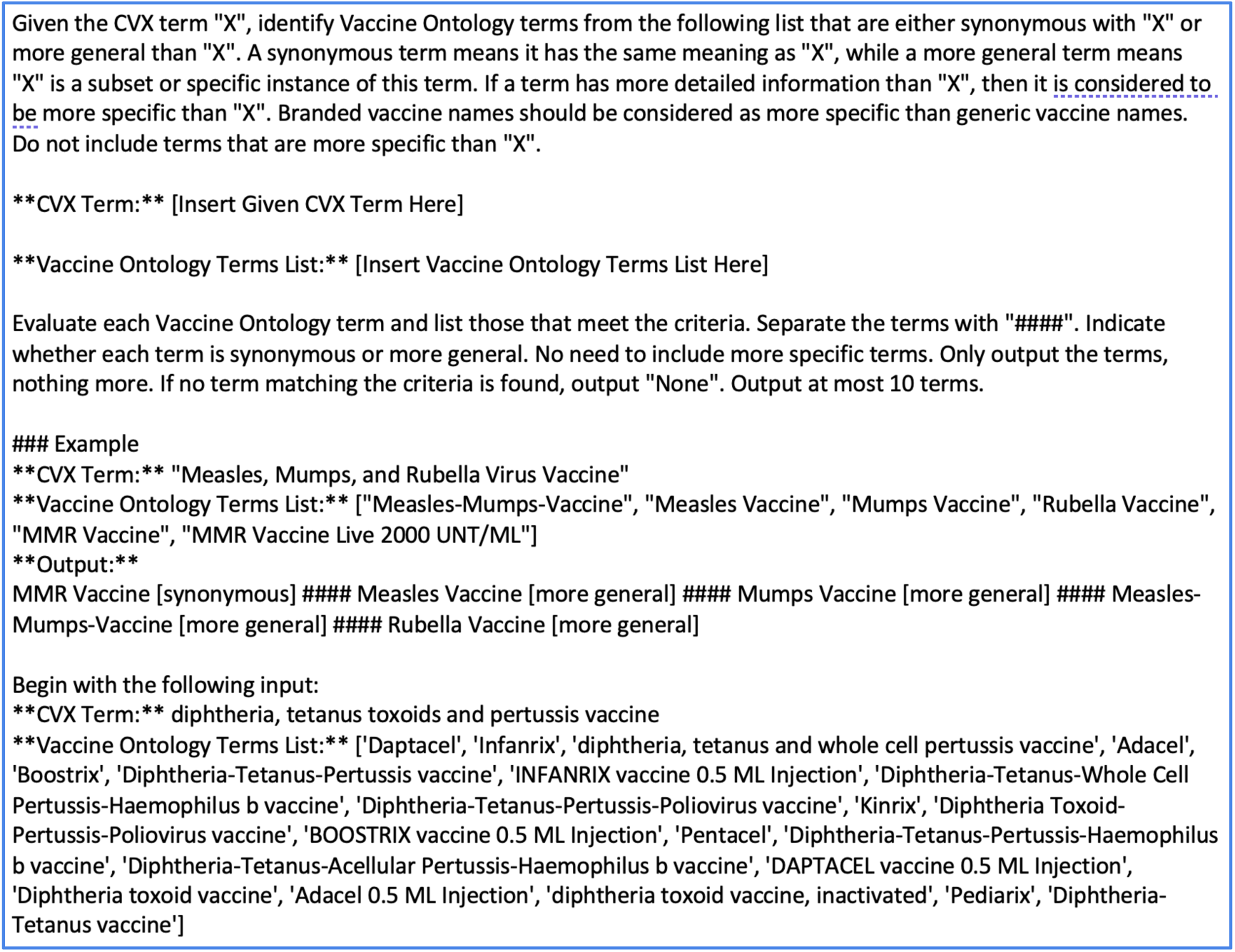
The prompt developed for GPT-4o to further refine the candidate mappings identified by the embedding-based approach.

#### 2.3.5. Evaluation of semi-automated CVX-VO mappings

Manual annotation was used to evaluate the semi-automated CVX-VO mapping results. As mentioned earlier, for each CVX term, the method suggested at most 10 VO candidate matches, or the method suggests that the CVX term is unmappable to VO and hence a new corresponding term would be needed to be added to VO. The suggestions made by our best-performing approach on the 2022-10-05 version were handed over to the curators for their review and potential adaptation to VO. For each CVX term, the validity of the suggestions was assessed. For a CVX term, if the approach suggested candidate mappings, and these contained the correct match, then this was considered as a valid case. On the other hand, if the approach does not suggest any candidate mappings (i.e. the CVX term is unmappable to VO), then it was checked whether there are any VO terms that the CVX term can be aligned with. If not, this was considered as a valid case as well.

### 2.4. Data access and license

After all the CVX terms are mapped to VO, a specific CVX-VO OWL file was generated and available at the CVX-VO GitHub website (https://github.com/vaccineontology/VO/tree/master/CVX-VO), which is under the general VO GitHub (https://github.com/vaccineontology/VO). The specific VO-to-CVX mapping SSSOM file is available at: https://github.com/vaccineontology/VO/blob/master/mappings/vo-cvx.sssom.tsv. The mapping file includes the CVX-to-VO code mappings, CVX vaccine full names and descriptions of VO. The OWL file of the CVX-VO view is also available at: http://purl.obolibrary.org/obo/vo/cvx-vo.owl. The CVX-VO mapping related data and code are freely available with the access license the same as the VO license, i.e., license CC BY 4.0 (http://creativecommons.org/licenses/by/4.0/).

## 3. Results

### 3.1. CVX classification

Current CVX vocabulary has 273 terms, including different kinds of vaccine terms and a small part of non-vaccine terms all presented in a code list without hierarchy. To better understand the CVX system, we systematically classified the CVX terms based on specific vaccine types. Note that most CVX codes are for specific vaccines. However, CVX also has grouping/classification codes such as ‘*polio, unspecified formulation*’ (CVX code: 89) and ‘*DTaP, unspecified formulation*’ (CVX code: 107). In our classification, we treated them as usual codes.

**Table 1** is a summary of the CVX terms as of May 22, 2024. Briefly, there are 273 terms in current version of CVX vocabulary, which can be classified to: 161 (58.9%) viral vaccines, 43 (15.7%) bacterial vaccines, 2 (0.7%) parasite vaccines, 33 (12.0%) combination vaccines, 1 (0.3%) cancer vaccine, 24 (8.7%) passive vaccines, and 9 other non-vaccine codes (**Table 1**, and the csv file in GitHub as indicated in Methods).

**Table 1.** CVX term classification and summary.

| Vaccine group | Vaccine types | Number of terms |
| --- | --- | --- |
| Virus vaccine (single) |  | **161 (58.9%)** |
|  | *Influenza vaccine* | 33 |
|  | *SARS-COV-2 related vaccine* | 47 |
|  | *Hepatitis vaccine* | 16 |
|  | *Other viral vaccines* | 65 |
| Bacterial vaccine (single) |  | **43 (15.7%)** |
|  | *Meningococcal vaccine* | 10 |
|  | *Pneumococcal vaccine* | 8 |
|  | *Other bacterial vaccine* | 25 |
| Parasite vaccine |  | **2 (0.7%)** |
|  | *Leishmaniasis vaccine* |  |
|  | *Malaria vaccine* |  |
| Combination Vaccines |  | **33 (12.0%)** |
|  | *Bacteria + bacteria* | 18 |
|  | *Virus + virus* | 5 |
|  | *Bacteria + virus* | 10 |
| Cancer vaccine |  | **1 (0.3%)** |
|  | *Melanoma vaccine* | 1 |
| Passive vaccine |  | **24 (8.7%)** |
| Other codes |  | **9 (3.6%)** |

The total 161 viral vaccines target 27 viral pathogens. These include 47 SARS-CoV-2 related vaccines, 33 influenza vaccines, and 16 hepatitis vaccines (**Table 1**). The remaining 65 vaccines target 24 different viral pathogens, such as 5 vaccines for rabies virus, 5 vaccines for human papillomavirus, and 4 vaccines for polio virus, among others.

The whole list of 43 bacterial vaccines recorded in CVX include 10 meningococcal vaccines, 8 pneumococcal vaccines, 6 typhoid vaccines, 4 cholera vaccines and 15 other bacterial vaccines. While ‘*Staphylococcus bacteriophage lysate*’ contains components of *Staphylococcus aureus* (*S. aureus*), a bacteriophage and some culture medium ingredients^21,22^, and it is designed to stimulate an immune response against staphylococcal infections, so it is considered as a type of bacterial vaccine in VO. A total of 33 combination vaccine terms exist in CVX. A large part of them are DTP (diphtheria, tetanus toxoids and pertussis) related combination vaccines. There are two parasite vaccines recorded in CVX: ‘*leishmaniasis vaccine*’ and ‘*malaria vaccine*’. There is only one cancer vaccine term in CVX: ‘*Melanoma vaccine*’.

There are 9 other codes that are not typical vaccine terms (**Table 2**). ‘*AS03 adjuvant*’ is a specific vaccine adjuvant. These codes also include 4 codes related to tuberculin skin test such as ‘*tuberculin skin test; unspecified formulation*’ and ‘*tuberculin skin test; purified protein derivative solution, intradermal*’, which are classified as diagnostic tests for tuberculosis. CVX also includes several terms that are not specific vaccines but are related to vaccines or vaccination, including ‘*Historical record of a typhus vaccination*’, ‘*no vaccine administered*’, *‘unknown vaccine and immune globulin’*, and ‘*RESERVED - do not use*’.

**Table 2.** Non-vaccine CVX codes.

| CVX term name | CVX code | VO code |
| --- | --- | --- |
| unknown vaccine or immune globulin | 999 | VO_0010150 |
| AS03 Adjuvant | 801 | VO_0001320 |
| no vaccine administered | 998 | VO_0010149 |
| RESERVED - do not use | 99 | VO_0010229 |
| tuberculin skin test; unspecified formulation | 98 | VO_0010233 |
| tuberculin skin test; old tuberculin, multipuncture device | 95 | VO_0010230 |
| tuberculin skin test; purified protein derivative solution, intradermal | 96 | VO_0010231 |
| tuberculin skin test; purified protein derivative, multipuncture device | 97 | VO_0010232 |
| Historical record of a typhus vaccination | 131 | VO_0006019 |

In addition, CVX includes 24 immune globulins, antibodies, and antitoxins. These terms are not considered as typical vaccines. However, they generate the immune responses and have been proposed to be passive vaccines^22^, which is further modeled and represented using the VO classification as detailed later in the paper.

### 3.2. VO high-level design for classification of CVX vaccine terms

Our basic CVX-to-VO mapping strategy is to reuse the VO hierarchy and map each CVX term to VO. Initially, we found that some CVX terms do not have exact VO term mapping and some CVX terms could be possibly mapped to multiple VO terms. After discussion, we generated and stuck with an “exact mapping” rule: Each CVX term is best mapped to one specific VO term and if there is no exact mapping, we will re-generate a new VO term that could be mapped to the CVX term.

**Figure 2** illustrates a high-level hierarchical design of CVX-mapped VO terms and their associated terms. Specifically, most of the current 273 CVX terms can be classified under the VO: vaccine branch. An exception is the CVX term ‘*AS03 adjuvant*’, which is classified under ‘*vaccine adjuvant*’ in VO (**Figure 2A**). The four tuberculin skin test terms are classified under OBCS:‘*diagnostic process*’ (OGMS_0000104). Several other CVX codes such as ‘*RESERVED - do not use*’ and ‘*Historical record of a typhus vaccination*’ are classified as ‘*information content entity*’ (IAO_0000030) in VO.

**Figure 2.**
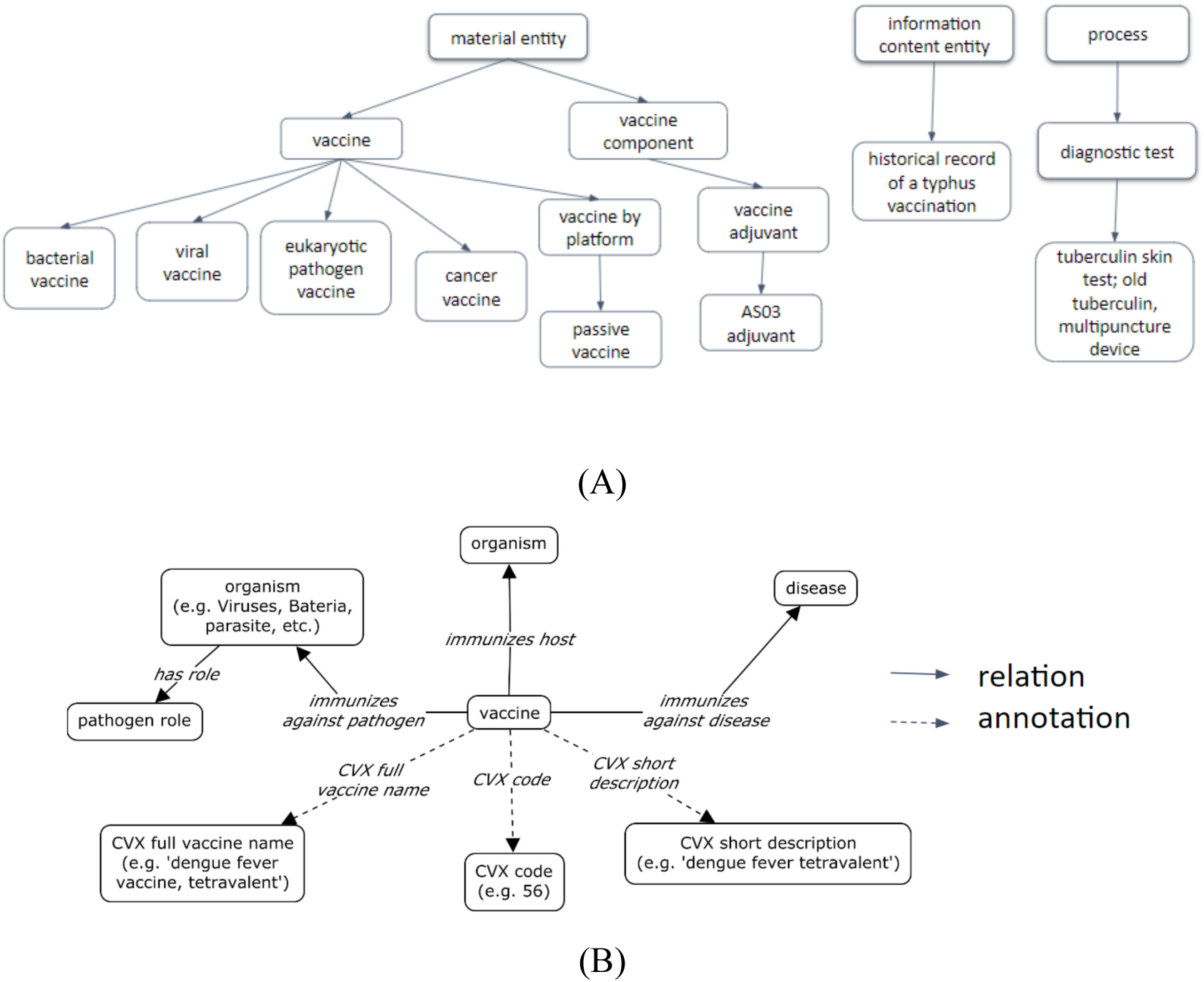
CVX-VO general ontology design. (A) high-level hierarchical VO structure that aligns with CVX. (B) CVX-VO relational design pattern of CVX-related terms.

Furthermore, we developed an ontology design pattern that represents the relations among different vaccine-related entities (**Figure 2B**). Specifically, in the VO setting, a vaccine ‘*immunizes host*’ some organism (e.g. human), ‘*immunizes against pathogen*’ some pathogen (e.g., influenza virus), and ‘*immunizes against disease*’ some disease (e.g., influenza). For each vaccine with specific CVX term mapping, we have added three specific annotation properties, including ‘*CVX full vaccine name*’ (e.g., dengue fever vaccine, tetravalent), ‘*CVX code*’ (e.g., 55), and ‘*CVX short description*’ (e.g., dengue fever tetravalent).

**Figure 3** demonstrates how the VO maps and represents a specific CVX term: ‘*SARS-COV-2 (COVID-19) vaccine, mRNA, spike protein, LNP, preservative free, 30 mcg/0.3mL dose*’ (CVX code: 208). This mapping was done by using annotation property “*CVX code: 208*”, ‘*CVX full name: SARS-COV-2 (COVID-19) vaccine, mRNA, spike protein, LNP, preservative free, 30 mcg/0.3mL dose*’, and ‘*CVX short description: COVID-19, mRNA, LNP-S,PF, 30 mcg/0.3mL dose*’ in VO to ensure a comprehensive integration of CVX contents. VO hierarchy was shown as ‘*SARS-COV-2 (COVID-19) vaccine, mRNA, spike protein, LNP, preservative free, 30 mcg/0.3mL dose*’ (VO_0006062) is under the parent term of ‘*authorized COVID-19 RNA vaccine*’ (VO_0005265), together with *‘COVID-19 vaccines*’ (VO_0004908), and their relations are under the hierarchy of taxonomy. Entities associated with this vaccine shown were implemented as logic axioms of a COVID-19 vaccine in VO.

**Figure 3.**
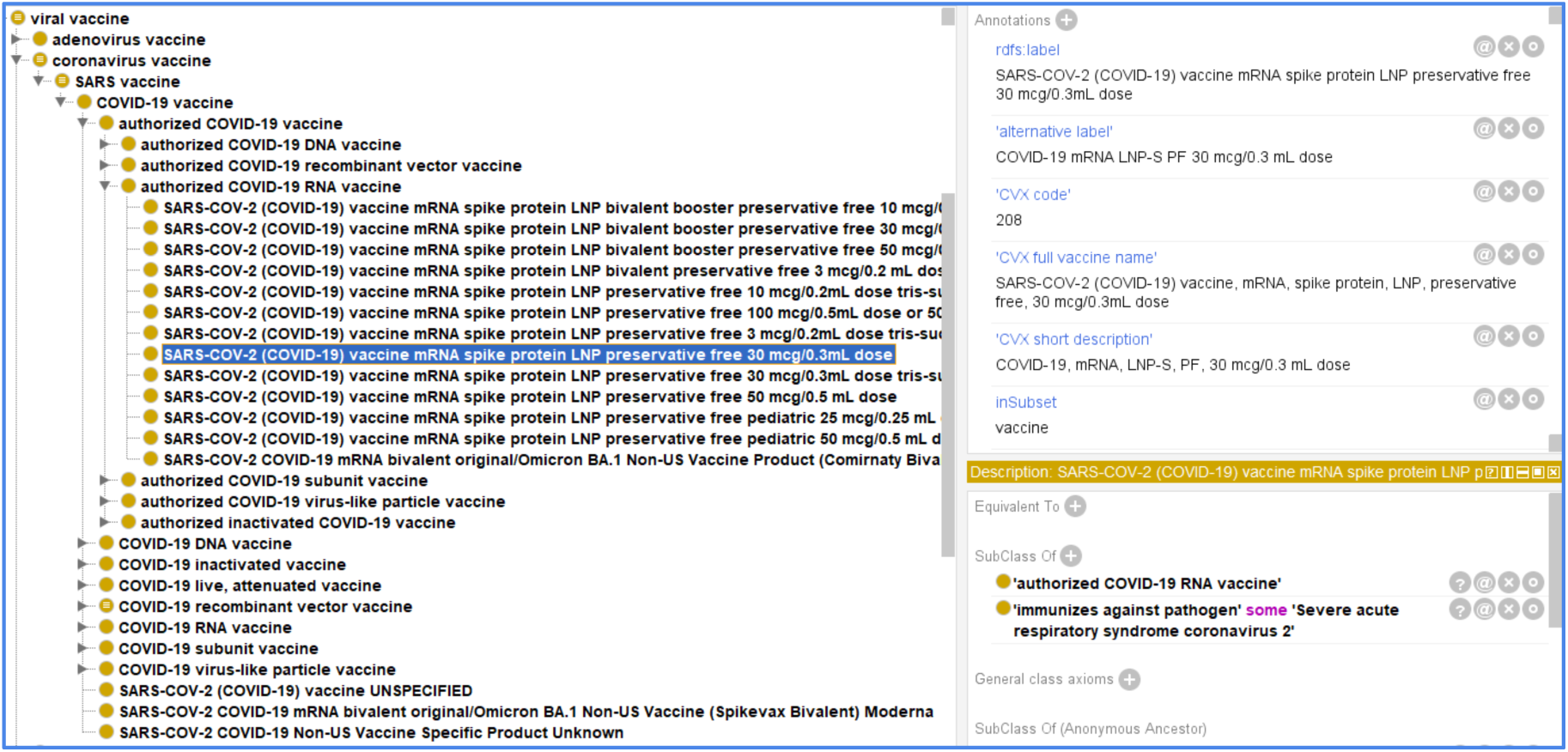
An example of VO term to illustrate how VO maps to CVX. Protégé-OWL editor was used for the ontology visualization and editing.

### 3.3. Semi-automated approaches for CVX-VO mapping

We applied our approaches on the 2022-10-05 release of VO, which contained 4,096 vaccine terms under the VO concept ‘*Vaccine*’ (VO_0000001). As shown in **Table 3**, the word-level similarity-based refinement approach, with an accuracy of 0.8555 produced the best performance. Out of the 263 CVX terms, mappings to VO were identified for 30, and missing concepts in VO were identified for 233. Based on the ascertainment of the curators, 21 out of 30 mappings as well as 204 out of 233 missing concepts in VO were valid, leading to a total of 225 out of the 263 suggestions being valid (an accuracy of 85.55%).

**Table 3.** Performance of each model in identifying CVX to VO mappings or missing VO concepts for the 2022-10-05 release of VO.

| Method | Predicted |  |  | Valid |  |  | Accuracy |
| --- | --- | --- | --- | --- | --- | --- | --- |
|  | Mapped | Missing | Total | Mapped | Missing | Total |  |
| Embedding Similarity | 131 | 132 | 263 | 31 | 116 | 147 | 0.5589 |
| Word-level similarity-based refinement | 30 | 233 | 263 | 21 | 204 | 225 | **0.8555** |
| Large Language Model-based refinement | 76 | 187 | 263 | 30 | 163 | 193 | 0.7338 |

Based on these suggestions and various other internal maintenance procedures, VO was iteratively improved and released on 2024-06-01 with mappings with all CVX terms. To further investigate the abilities of our method, we applied it to the 2024-06-01 release and examined the suggestions made for the 263 CVX terms. The word-level similarity-based refinement approach suggested VO mappings for 229 CVX terms and also suggested 34 missing VO concepts. It was seen that 215 of our 229 CVX to VO mappings were valid. Since this version of VO contains mapped concepts for all CVX terms, no concept is missing; hence, all the missing concept suggestions were invalid. Therefore, altogether, 229 out of 263 suggestions made on the 2024-06-01 version were found to be valid (accuracy of 87.07%).

To demonstrate the CVX to VO mapping, **Table 4** shows four representative value CVX to VO mappings as identified by the word-level similarity-based refinement approach.

**Table 4.** Five valid CVX to VO mappings identified by the word-level similarity-based refinement approach for the 2024-06-01 release of VO.

| CVX Term | VO Term |
| --- | --- |
| CVX_29: cytomegalovirus immune globulin, intravenous | VO_0005441: cytomegalovirus immune globulin passive vaccine, intravenous |
| CVX_166: influenza, intradermal, quadrivalent, preservative free, injectable | VO_0005463: Influenza, injectable, MDCK, quadrivalent, preservative |
| CVX_300: SARS-COV-2 (COVID-19) vaccine, mRNA, spike protein, LNP, bivalent, preservative free, 30 mcg/0.3 mL dose, tris-sucrose formulation | VO_0006056: SARS-COV-2 (COVID-19) vaccine mRNA spike protein LNP bivalent booster preservative free 30 mcg/0.3 mL dose tris-sucrose formulation |
| CVX_156: Rho(D) Immune globulin- IV or IM | VO_0005450: Rho(D) Immune globulin- IV or IM passive vaccine |
| CVX_186: Influenza, injectable, Madin Darby Canine Kidney, quadrivalent with preservative | VO_0005458: Influenza, injectable, Madin Darby Canine Kidney, quadrivalent with preservative |

### 3.4. Evaluation and improvement of the VO by identifying missing terms

The CVX-VO mapping and VO updatings were completed by a combined usage of manual and semi-automated mapping. The semi-automatic mapping methods were improved over time, together with our dynamic manual evaluation process. The manual and semi-automatic mapping teams learned from each other during the whole dynamic mapping process.

One lesson learned in the mapping process was the change of mapping strategy. Initially the one-to-many strategy was used, and later we changed the mapping strategy to one-to-one exact mappings in order to ensure the high quality mapping. Our research found that many individual CVX codes could be mapped to more than one VO terms that are parent and child. For example, CVX code 03 ‘*measles, mumps and rubella virus vaccine*’ mapped to: ‘*Measles-Mumps-Rubella vaccine*’ (VO_0000731), and ‘*M-M-R II*’ (VO_0000069). However, our examination found that the ‘*Measles-Mumps-Rubella vaccine*’ (VO_0000731) is an exact mapping with the CVX code and the M-M-R II is a specific vaccine brand name that is classified as a subclass under the VO_0000731. Based on our exact mapping rule, we only mapped the CVX code to the VO_0000731 but not VO_0000069. However, since the VO asserts the branded vaccine M-M-R II under ‘*Measles-Mumps-Rubella vaccine*’, M-M-R II could be easily inferred as a specific type of the CVX vaccine.

### 3.5. Proposing “passive vaccine” in VO and its usage for CVX mapping

Notably, CVX includes 24 immune globulin, antibody and antitoxin terms. Our community-based research and discussion eventually achieved a consensus that these should be classified as passive vaccines in VO. According to VO, passive vaccine is defined as “a vaccine that is made of immune globulin (and antitoxin) that is used to immunize an individual against some disease”^24^. It is “passive” because instead of having the host organism actively producing the immune product (e.g., immune globulin) after active vaccination, the host is passively given the immune product that is directly used against disease without complex immune stimulation processes in vivo. Passive vaccine is the entity that is used in the process of passive immunization, while passive immunization is defined as the direct administration of preformed immune globulins and antitoxin to an individual^25^. Many studies have classified disease-specific antibodies (i.e. immune globulins) as ‘passive vaccines’^26–29^. The classification of them as passive vaccines is important for semantic integration and analysis of vaccine-related terms. Furthermore, it helps to enhance data consistency and interoperability of immunization processes with better alignment with current immunological frameworks.

As a result, 24 immune globulin terms were added as subclasses under ‘*passive vaccine*’ (VO_0005439) in the VO (**Table 5**). Proper hierarchies and annotations were also added.

**Table 5.**
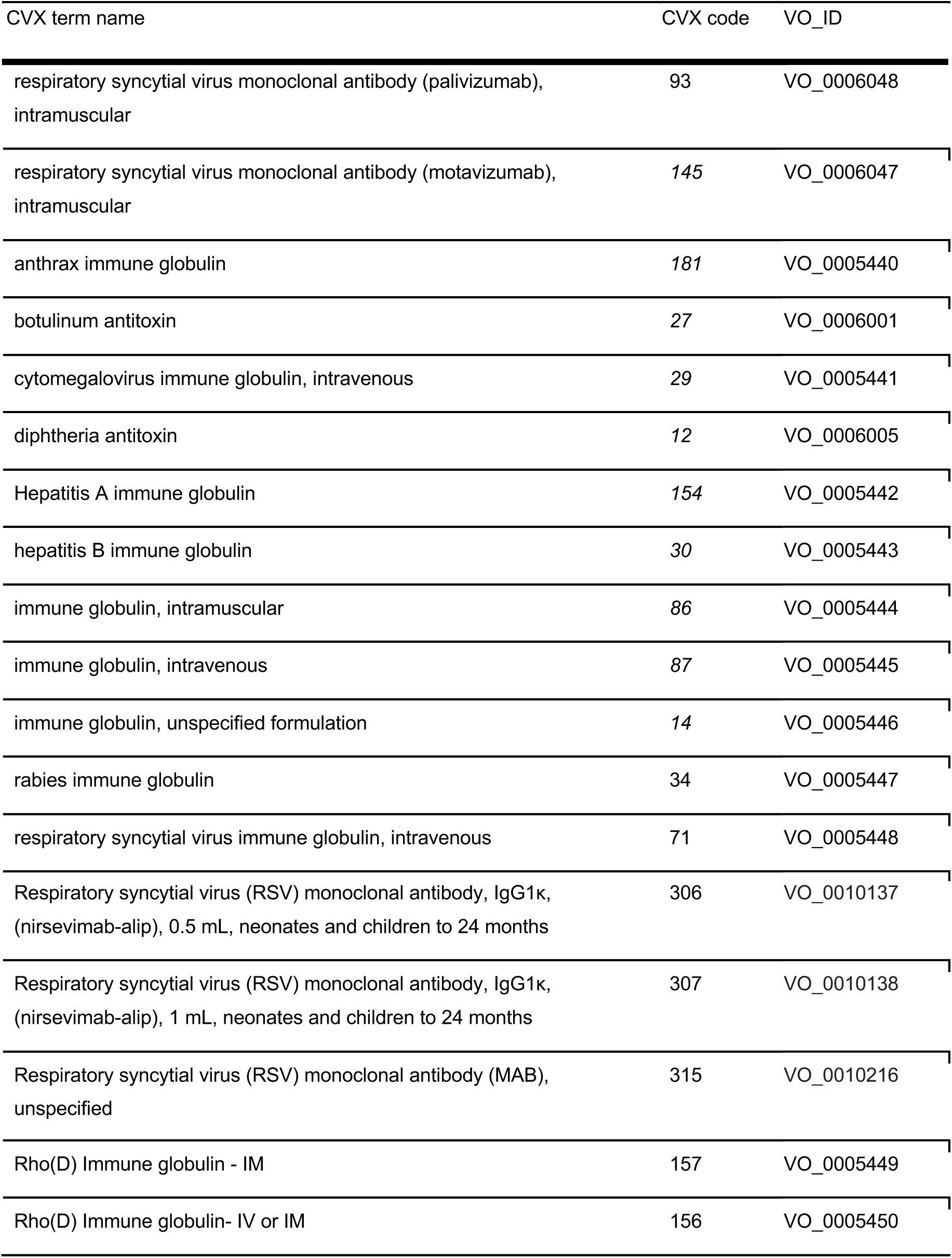
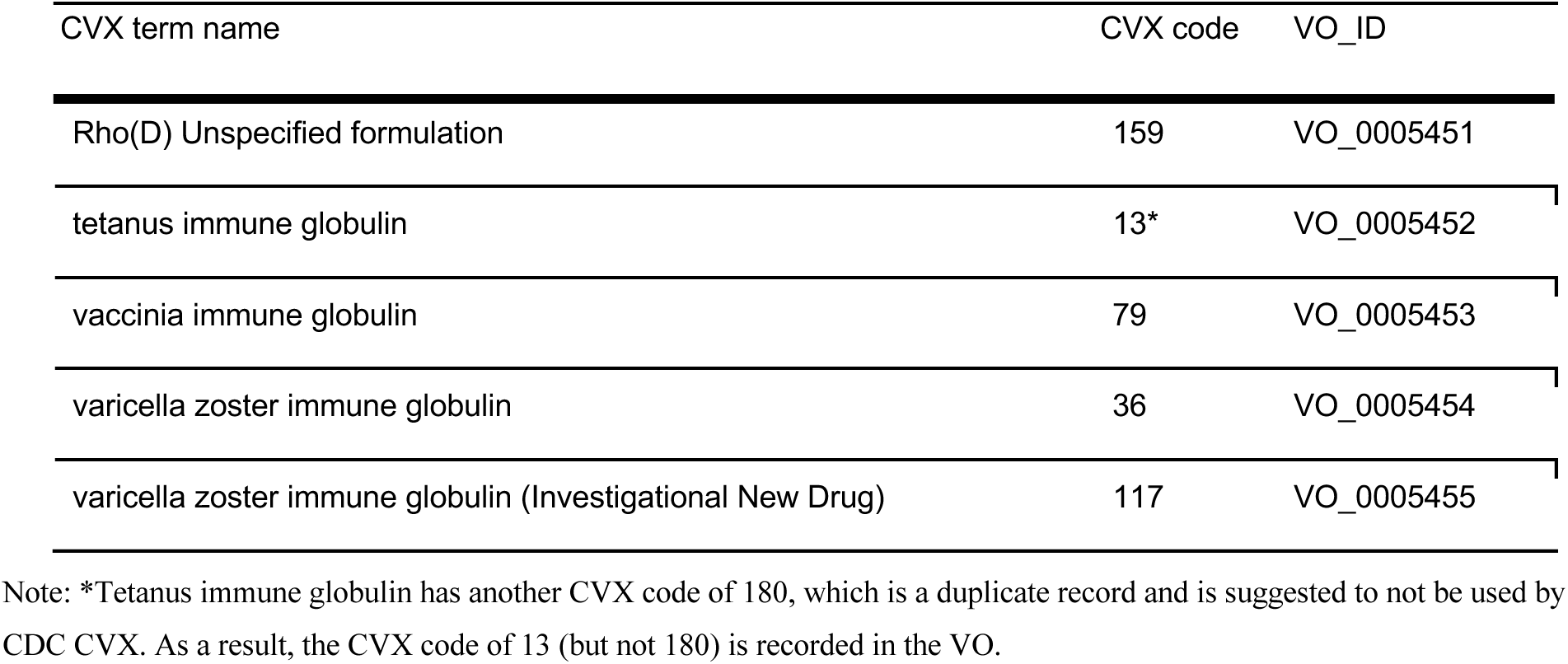
‘passive vaccines’ as recorded by CVX and VO.

| CVX term name | CVX code | VO_ID |
| --- | --- | --- |
| respiratory syncytial virus monoclonal antibody (palivizumab), intramuscular | 93 | VO_0006048 |
| respiratory syncytial virus monoclonal antibody (motavizumab), intramuscular | 145 | VO_0006047 |
| anthrax immune globulin | 181 | VO_0005440 |
| botulinum antitoxin | 27 | VO_0006001 |
| cytomegalovirus immune globulin, intravenous | 29 | VO_0005441 |
| diphtheria antitoxin | 12 | VO_0006005 |
| Hepatitis A immune globulin | 154 | VO_0005442 |
| hepatitis B immune globulin | 30 | VO_0005443 |
| immune globulin, intramuscular | 86 | VO_0005444 |
| immune globulin, intravenous | 87 | VO_0005445 |
| immune globulin, unspecified formulation | 14 | VO_0005446 |
| rabies immune globulin | 34 | VO_0005447 |
| respiratory syncytial virus immune globulin, intravenous | 71 | VO_0005448 |
| Respiratory syncytial virus (RSV) monoclonal antibody, IgG1k, (nirsevimab-alip), 0.5 mL, neonates and children to 24 months | 306 | VO_0010137 |
| Respiratory syncytial virus (RSV) monoclonal antibody, IgG1k, (nirsevimab-alip), 1 mL, neonates and children to 24 months | 307 | VO_0010138 |
| Respiratory syncytial virus (RSV) monoclonal antibody (MAB), unspecified | 315 | VO_0010216 |
| Rho(D) Immune globulin - IM | 157 | VO_0005449 |
| Rho(D) Immune globulin- IV or IM | 156 | VO_0005450 |
| Rho(D) Unspecified formulation | 159 | VO_0005451 |
| tetanus immune globulin | 13* | VO_0005452 |
| vaccinia immune globulin | 79 | VO_0005453 |
| varicella zoster immune globulin | 36 | VO_0005454 |
| varicella zoster immune globulin (Investigational New Drug) | 117 | VO_0005455 |

Note: \*Tetanus immune globulin has another CVX code of 180, which is a duplicate record and is suggested to not be used by CDC CVX. As a result, the CVX code of 13 (but not 180) is recorded in the VO.

### 3.6. Statistics of CVX-to-VO mapping and OWL generation

Overall, out of 273 vaccine terms in CVX, a total of 273 CVX-VO mapping pairs were identified or generated, which means that every CVX term has its specific VO mapping. The identified mapping information is recorded in a csv file as indicated in the Methods section.

The manually annotated file was used to support the development and evaluation of the automatic mapping methods as indicated above. Meanwhile, the automatic mapping algorithms developed significantly helped the VO mapping and classification. It is also noted that the latest 2024-06-01 release is a significant improvement over the earlier 2022-10-05 release of VO.

CVX vaccines constitute a small portion of vaccines in VO. To facilitate working with a CVX-VO mapped vaccines only ontology, we generated the CVX-VO view (cvx-vo.owl), as indicated in the Methods section, by retrieving all CVX-VO mapped vaccines terms and associated axioms from the full version VO. To add more semantic relations, the cvx-vo.owl provides additional terms that show the hierarchical terms and relations among the CVX-mapped terms.

### 3.7. Use cases of CVX-VO mapping: Computer-assisted queries of vaccine groups based on CVX-VO hierarchies

The CVX-VO mapping can be used to support different applications, such as automatic computer-assisted queries of vaccine groups. As a demonstration, **Figure 4** shows a description logic (DL)-query that queries for how many H1N1 influenza vaccines recorded in the CVX-VO mapping OWL file. The VO represents the H1N1 influenza vaccine using the following axiom:

> ‘immunizes against pathogen’ *some* ‘H1N1 subtype’.In total, CVX records four specific H1N1 influenza, which can be clearly identified using our DL-query (**Figure 4**). Similarly, we can also use SPARQL to perform such a query (data not shown). Such simple queries empowered by our VO mapping illustrates how we can use semantics-based ontology to support CVX ontologization and various use cases.

**Figure 4.**
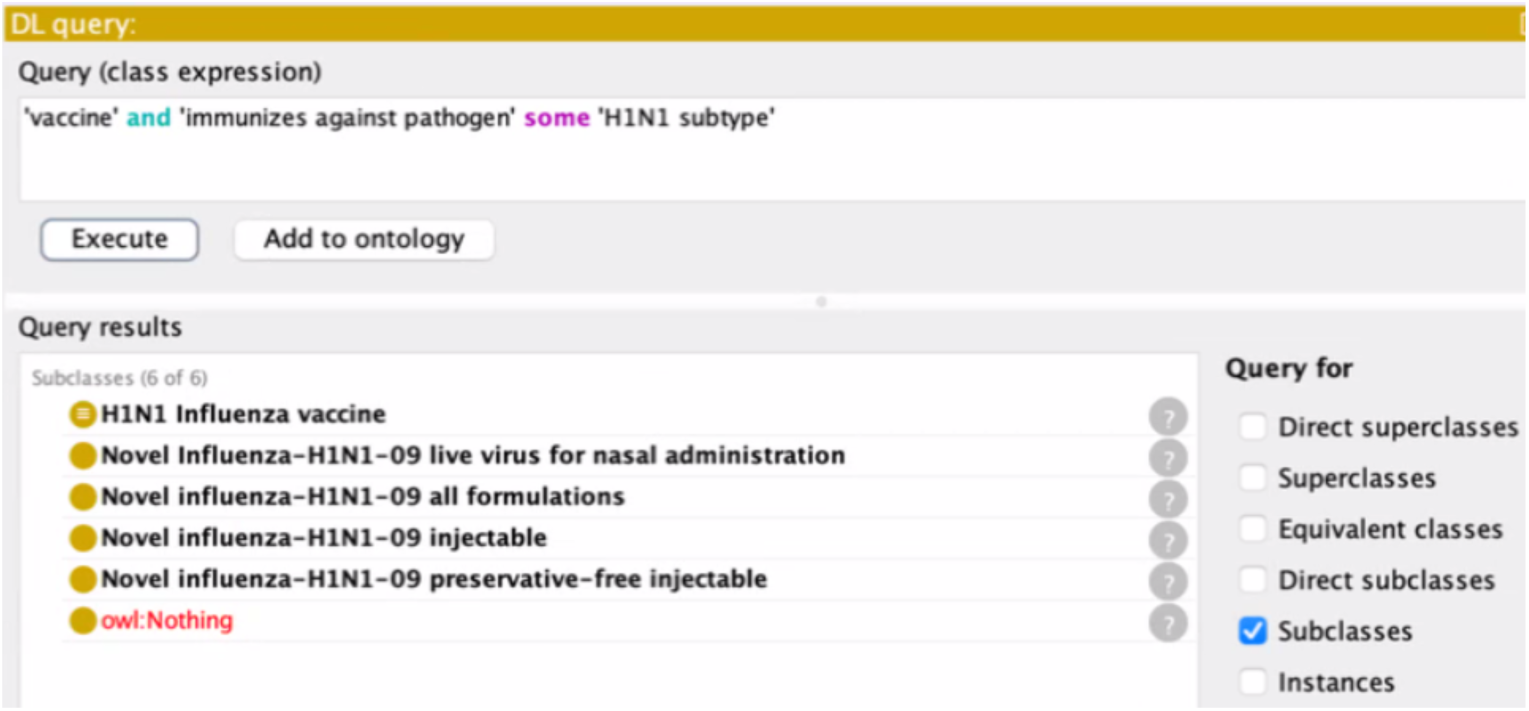
DL-query of H1N1 influenza vaccines.

In addition, we were also able to develop and apply DL queries or SPARQL queries to directly query the Vaccine Ontology (VO) (instead of the CVX-VO mapping OWL file). The VO-based queries were able to identify more vaccines not mapped to CVX. For example, by querying VO for ‘*Measles-Mumps-Rubella vaccine*’ (VO_0000731) (query scripts not shown), we were able to find many vaccines with brand names under this vaccine branch, such as ‘*M-M-R II*’ (VO_0000069), ‘*Morupar*’ (VO_0001013), and ‘*Virivac*’ (VO_0003117).

## 4. Discussion

The contributions of this manuscript are multiple. First, we finished our CVX-to-VO mapping and alignment and generated the CVX-to-VO mapping OWL file. Second, we developed computational methods to automate the mapping pipeline for the CVX-to-VO mapping, and such methods were used for more efficient CVX to VO mappings. In addition, the mapping methods can and will also be used for later mapping between VO and other OMOP vaccine vocabularies such as RxNorm and RxNorm Extension. Third, we have provided design patterns and use case demonstrations to show the value of the VO mapping to support better representation of terms in the OMOP ontology system based on the OHDSI’s OMOP Common Data Model. We innovatively proposed the inclusion of the ‘passive vaccine’ branch in VO that includes 24 immunoglobulins and antitoxins from CVX as passive vaccines. One specific use case is computer-assisted queries of vaccine groups based on the CVX-VO hierarchies. It is also possible to extend the query to identify vaccines with brand names that match to the CVX vaccines.

### 4.1. Semi-automated approaches for CVX-VO mapping

Overall, we applied both manual and semi-automated methods to map CVX and VO vaccine terms and updated VO correspondingly. The semi-automated methods can be promising as they require significantly less human effort than purely manual approaches.

It should be noted that while the manual mapping of CVX terms to VO might not warrant the development of an automated method, our semi-automatic method assisted the manual curation process by providing either VO candidate concepts or the suggestion that a new concept should be added to VO. Furthermore, this method will serve as the baseline for matching and evaluating the process of integrating terms from other larger vocabularies such as RxNorm and RxNorm Extension. Our plan is to use the VO as an integrative platform to systematically represent the mapped results for vaccine terms in individual vocabularies. We started with the mapping of CVX vaccine terms to the VO using manual and semi-automatic strategies.

For a given CVX term, our semi-automated approaches suggest either candidate concepts from the Vaccine Ontology that are likely to be mapped or indicate that the CVX term is unmappable to current VO terms, suggesting a new concept addition to VO. The suggestions were handed over to the curators for review and potential incorporation into VO in a new release. We have investigated three such semi-automated methods: a semantic approach in the form of sentence embedding using pre-trained transformer models, two hybrid approaches using sentence embeddings as the initial matcher, and either 1) a purely lexical approach with Jaccard similarity comparison or 2) Large Language Model (LLM)-based approach as the final matcher. Of the three approaches, the first hybrid approach was found to be the best in terms of the validity of its suggestions.

It should be noted that the word-level similarity-based approach achieved the best performance from the implemented approaches outperforming the LLM-based approach. However the LLM used here was not fine-tuned or pre-trained for this specific task and may not have had sufficient training to distinguish between related but nuanced vaccine concepts while the word-level similarity based approach is agnostic to such nuances.

While the word-level similarity-based refinement approach Hybrid approach 1 outperforms the other matching approaches, it would be interesting to evaluate sentence embedding and LLM prompt engineering with models that are fine-tuned or pre-trained on a more specific biomedical domain. Furthermore, it would be more impactful to the curation process if the automated approach is extended to provide concept names for the new concepts needing to be added to VO. In addition, if such an approach could automatically identify logical definitions (attributes) of those new concepts, this would significantly reduce the curation time. We also intend to extend the application of the semi-automated approaches to other external vocabulary systems to contribute to enhanced vaccine data integration within VO. While the lexical and LLM approaches are highly task-specific and would require further analysis and evaluation prior to implementation, such approaches are especially important when dealing with large external terminologies like RxNorm and therefore will come in handy in supporting the manual effort of the curators. With expanded coverage and interoperability, the updated VO will further be used for systematic and integrative analysis of vaccine-related clinical data available in the OHDSI/OMOP compliant systems.

### 4.2. Manual mapping and VO updating

The CVX-to-VO mapping offers many advantages. First, such mapping converts the full list of CVX terms to the ontological representation in the vaccine ontology (VO), making us have a better understanding of the hierarchical structure and relations among these terms. Second, since the VO also represents many other types of vaccines at different stages (including licensed, clinical trials, and research stage), the CVX-to-VO mapping allows us to interlink various types of vaccines defined in CVX to other vaccine terms defined in the VO. Third, such mapping also enables the development of many tools and programs. As a community standard, VO has been widely used in different systems such ImmPort^30^, Human Immunology Project Consortium (HIPC)^31^, and Vaccine Adjuvant Compendium (VAC)^32^. Such mapping allows the seamless integration of different types of vaccine data.

One significant contribution of this study is the definition and enrichment of the ‘passive vaccine’ branch in VO and the CVX-to-VO mapping file. Notably, CVX includes 24 immune globulin vaccines, antibody and antitoxin terms, which were initially all missed in VO before this research. After group discussion, we formed a consensus that these should be classified as passive vaccines. All these immune globulin terms were then added as subclasses under ‘immune globulin passive vaccine’, a subclass of ‘passive vaccine’ in the VO. Proper hierarchies and annotations were also added.

### 4.3. Future work

The work of CVX to VO mapping and VO improvement well prepares us to proceed further for more complex mapping to other OMOP CDM vaccine containing terminologies, including RxNorm and RxNorm Extension. This is a team effort which will be performed by the members of the OHDSI Vaccine Vocabulary Working Group (Vaccine Vocab WG). We expect to incorporate the VO into the OHDSI Standardized vocabularies ontology system^33^ to enhance vaccine data standardization and interoperability in the large-scale network studies around the world. Such work will significantly enhance the coverage of the VO and its wide application.

## 5. Conclusion

In this study, we have done mapping and harmonization of CVX vaccine terms to the Vaccine Ontology, demonstrating that created CVX-to-VO mappings provide better semantic classification and applications, further supporting vaccine observational research on the real-world data such as EHR. We also investigated semi-automated approaches with good performance to facilitate future mapping work. The semi-automated approaches either suggest a list of candidate VO mapping for an external vocabulary term or indicate that the external vocabulary term is unmappable to VO requiring a new term to be added to VO. We investigated three approaches: (1) Embedding-based approach; (2) Hybrid approach 1 refining embedding-based baseline with word-level similarity; and (3) Hybrid approach 2 refining embedding-based baseline with a Large Language Model. The Hybrid approach 1 was found to be the best performing approach with an accuracy of 85.55% when applied on the 2022-10-05 release of VO.

## 6. Declarations

## Ethics approval

Not applicable.

## Consent to participate

Not applicable.

## Consent for publication

Not applicable.

## Competing interests

The authors declare no competing interests.

## Availability of data and materials

CVX-VO OWL file is available at the CVX-VO GitHub website (https://github.com/vaccineontology/VO/tree/master/CVX-VO), which is under the general VO GitHub (https://github.com/vaccineontology/VO). The specific VO-to-CVX mapping SSSOM file is available at: https://github.com/vaccineontology/VO/blob/master/mappings/vo-cvx.sssom.tsv. The CVX-VO mapping related data and code are freely available with the access license the same as the VO license, i.e., license CC BY 4.0 (http://creativecommons.org/licenses/by/4.0/).

## Funding

This study is supported by the National Institutes of Health (NIH) through grants U24AI171008 and R01NS116287, as well as the National Science Foundation (NSF) through grant 2047001. The content is solely the responsibility of the authors and does not necessarily represent the official views of the NIH or NSF.

## Author Contributions

YP conducted and completed the manual CVX to VO mappings. WM were responsible for semi-automated approaches for mapping CVX to VO. RA engineered the prompt for the Hybrid approach 2. YH and JZ were responsible for the ontology design pattern. JZ generated new terms into VO as an ontology expert. AD and QY served as domain experts of OHDSI vaccine classification. AD, QY, AYL and XY served as clinical vaccine domain experts. YH and LC conceptualized and supervised the whole study. YP, YH, WM and RA wrote the manuscript and generated figures and tables. All authors contributed to reviewing and revising the paper. All authors reviewed and approved of submission.

## Acknowledgements

We acknowledge the discussion and suggestions from the OHDSI community.

## Abbreviation

EHR: electronic health records
OMOP CDM: Observational Medical Outcomes Partnership Common Data Model
CVX: CDC Vaccine Administered code
HL7: Health Level Seven International
VO: Vaccine Ontology
OHDSI: Observational Health Data Sciences and Informatics
OBO: Open Biological and Biomedical Ontology
NCIRD: National Center of Immunization and Respiratory Diseases
CDC: Centers for Disease Control and Prevention
VAERS: Vaccine Adverse Event Analysis System
HIPC: Human Immunology Project Consortium
VAC: Vaccine Adjuvant Compendium
Vaccine Vocab WG: Vaccine Vocabulary Working Group

## Notes

https://github.com/vaccineontology/VO/tree/master/CVX-VO

http://purl.obolibrary.org/obo/vo/cvx-vo.owl

https://github.com/vaccineontology/VO/blob/master/mappings/vo-cvx.sssom.tsv

https://github.com/vaccineontology/VO

### Competing Interest Statement

The authors have declared no competing interest.

